# Additional Essential Oils with High Activity against Stationary Phase *Borrelia burgdorferi*

**DOI:** 10.1101/260091

**Authors:** Jie Feng, Wanliang Shi, Judith Miklossy, Ying Zhang

## Abstract

Lyme disease is the most common vector borne-disease in the US. While the majority of the Lyme disease patients can be cured with 2-4 week antibiotic treatment, about 10-20% of patients continue to suffer from persisting symptoms. While the cause of this condition is unclear, persistent infection was proposed as one possibility. It has recently been shown that *B. burgdorferi* develops dormant persisters in stationary phase cultures that are not killed by the current Lyme antibiotics, and there is interest to identify novel drug candidates that more effectively kill such forms. We previously evaluated 34 essential oils and identified some highly active candidates with excellent activity against biofilm and stationary phase *B. burgdorferi.* Here we screened another 35 essential oils and found 10 essential oils (garlic, allspice, cumin, palmarosa, myrrh, hedycheim, amyris, thyme white, litsea cubeba, lemon eucalyptus) and the active component of cinnamon bark cinnamaldehyde (CA) at a low concentration of 0.1% to have high activity against stationary phase *B. burgdorferi.* At a very low 0.05% concentration, garlic, allspice, palmarosa and CA still exhibited strong activity against the stationary phase *B. burgdorferi*. CA also showed strong activity against replicating *B. burgdorferi*, with a MIC of 0.02% (or 0.2 μg/mL). In subculture studies, the top 5 hits garlic, allspice, myrrh, hedycheim, and litsea cubeba completely eradicated all *B. burgdorferi* stationary phase cells at 0.1%, while palmarosa, lemon eucalyptus, amyris, cumin, and thyme white failed to do so as shown by visible spirochetal growth after 21-day subculture. At 0.05% concentration, only garlic essential oil and CA sterilized the *B. burgdorferi* stationary phase culture as shown by no regrowth during subculture, while allspice, myrrh, hedycheim and litsea cubeba all had visible growth during subculture. Future studies are needed to determine if these highly active essential oils could eradicate persistent *B. burgdorferi* infection in vivo.

## INTRODUCTION

Lyme disease, caused by the spirochetal organism *Borrelia burgdorferi*, is the most common vector borne-disease in the US with about 300,000 cases a year. While the majority of the Lyme disease patients can be cured with the standard 2-4 week antibiotic monotherapy with doxycycline or amoxicillin or cefuroxime, about 10-20% of patients continue to suffer from persisting symptoms of fatigue, joint or musculoskelital pain, and neuropsychiatric symptoms even 6 months after taking the standard antibiotic therapy. These latter patients suffer from a poorly understood condition, called post-treatment Lyme disease (PTLDS) syndrome. While the cause for this condition is unclear and is likely multifactorial, the following factors have been proposed to be involved: including autoimmunity, host response to dead debris of borrelia organism, tissue damage caused during the infection, and persistent infection. There have been various anecdotal reports demonstrating persistence of the organism despite standard antibiotic treatment. For example, culture of *B. burgdorferi* bacteria from patients despite treatment has been reported as infrequent case reports (Marques et al., 2014). In addition, in animal studies with mice, dogs and monkeys, it has been shown that the current Lyme antibiotic treatment with doxycycline, cefuroxime or ceftriaxone is unable to completely eradicate the borrelia organism as detected by xenodiagnosis and PCR but viable organism cannot be cultured in the conventional sense as in other persistent bacterial infections like tuberculosis after treatment (Zhang, 2004;Zhang et al., 2012).

Recently, it has been demonstrated that *B. burgdorferi* can form various dormant non-growing persisters in stationary phase cultures that are tolerant or not killed by the current antibiotics used to treat Lyme disease (Feng et al., 2014a;Caskey and Embers, 2015;Feng et al., 2015a;Sharma et al., 2015). Thus, while the current Lyme antibiotics are good at killing the growing *B. burgdorferi* they have poor activity against the non-growing persisters enriched in stationary phase culture (Feng et al., 2014a;Feng et al., 2015a;Feng et al., 2016). Therefore, there is interest to identify drugs that are more active against the *B. burgdorferi* persisters than the current Lyme antibiotics. We used the stationary phase culture of *B. burgdorferi* as a persister model and performed high throughput screens and identified a range of drug candidates such as daptomycin, clofazimine, sulfa drugs, daunomycin, etc. that have high activity against *B. burgdorferi* persisters. These persister active drugs act differently from the current Lyme antibiotics as they seem to preferentially target the membrane. We found that the variant persister forms such as round bodies, microcolonies, and biofilms with increasing degree of persistence in vitro, cannot be killed by the current Lyme antibiotics or even persister drugs like daptomycin alone, and that they can only be killed by a combination of drugs that kill persisters and drugs that kill the growing forms (Feng et al., 2015a). These observations provide a possible explanation in support of persistent infection despite antibiotic treatment in vivo.

Although daptomycin has good anti-persister activity, it is expensive and is an intravenous drug and difficult to administer and adopt in clinical setting and has limited penetration through blood brain barrier (BBB). Thus, there is interest to identify alternative drug candidates with high anti-persister activity. We recently screened a panel of 34 essential oils and found the top three candidates oregano oil and its active component carvacrol, cinnamon bark, and clove bud as having even better anti-persister activity than daptomycin at 40 uM (Feng et al., 2017). To identify more essential oils with high activity against *B. burgdorferi* persisters, in this study, we screened additional 35 different essential oils and found 10 essential oils (garlic, allspice, cumin, palmarosa, myrrh, hedycheim, amyris, thyme white, litsea cubeba, lemon eucalyptus) and the active compoent of cinnamon bark cinnamaldehyde as having strong activity in the stationary phase *B. burgdorferi* persister model.

## MATERIALS AND METHODS

### Strain, media and culture techniques

A low passaged *B. burgdorferi* strain B31 5A19 was kindly provided by Dr. Monica Embers (Caskey and Embers, 2015). The *B. burgdorferi* B31 strain was cultured in BSK-H medium (HiMedia Laboratories Pvt. Ltd.) with 6% rabbit serum (Sigma-Aldrich, St. Louis, MO, USA) in microaerophilic incubator (33°C, 5% CO2) without antibiotics. After incubation for 7 days, the stationary phase *B. burgdorferi* culture (~10^7^ spirochetes/mL) was transferred to a 96-well plate for evaluation of potential anti-persister activity of essential oils (see below).

### Essential oils and drugs

A panel of essential oils was purchased from Plant Guru (NJ, USA). Carvacrol, p-cymene, and α-terpinene were purchased from Sigma-Aldrich (USA). DMSO-soluble essential oils were dissolved in dimethyl sulfoxide (DMSO) at 10%, followed by dilution at 1:50 into 7-day old stationary phase cultures to achieve 0.2% final concentration. To make further dilutions for evaluating anti-borrelia activity, the 0.2% essential oils were further diluted with the stationary phase culture to achieve desired dilutions. DMSO-insoluble essential oils were added to *B. burgdorferi* cultures to form aqueous suspension by adequate vortexing, followed by immediate transfer of essential oil aqueous suspension in serial dilutions to desired concentrations and then added to *B. burgdorferi* cultures. Doxycycline (Dox), cefuroxime (CefU), (Sigma-Aldrich, USA) and daptomycin (Dap) (AK Scientific, Inc., USA) were dissolved in suitable solvents (The United States Pharmacopeial Convention, 2000;Wikler and National Committee for Clinical Laboratory, 2005) to form 5 mg/ml stock solutions. The antibiotic stocks were filter-sterilized by 0.2 μm filter and stored at -20°C.

### Microscopy

Treated *B. burgdorferi* cell suspensions were examined using BZ-X710 All-in-One fluorescence microscope (KEYENCE, Inc.). The SYBR Green I/PI viability assay was performed to assess the bacterial viability using the ratio of green/red fluorescence to determine the live:dead cell ratio, respectively, as described previously (Feng et al., 2014a;Feng et al., 2014b). This residual cell viability reading was confirmed by analyzing three representative images of the same bacterial cell suspension using fluorescence microscopy. BZ-X Analyzer and Image Pro-Plus software were used to quantitatively determine the fluorescence intensity.

### Evaluation of essential oils for their activities against *B. burgdorferi* stationary phase cultures

To evaluate the essential oils for possible activity against stationary phase *B. burgdorferi*, 4 μL DMSO stocks or aliquots of the essential oils were added to 96-well plate containing 100 μL of the 7-day old stationary phase *B. burgdorferi* culture to obtain the desired concentrations. In the primary essential oil screen, each essential oil was assayed in four concentrations, 1%, 0.5%, 0.25% and 0.125% (v/v) in 96-well plates. Daptomycin, doxycycline and cefuroxime were used as control drugs at 40 μM, 20 μM, 10 μM and 5 μM. The active hits were further confirmed with lower 0.1% and 0.05% concentrations; all tests were run in triplicate. All the plates were incubated at 33°C and 5% CO2 without shaking for 7 days when the residual viable cells remaining were measured using the SYBR Green I/PI viability assay and fluorescence microscopy as described (Feng et al., 2014a;Feng et al., 2014b).

### Antibiotic susceptibility testing

To qualitatively determine the effect of essential oils in a high-throughput manner, 4 μl of each essential oil from the pre-diluted stock was added to 100 μl 7-day old stationary phase *B. burgdorferi* culture in the 96-well plate. Plates were sealed and placed in 33°C microaerophilic incubator. After 7 day treatment the SYBR Green I/PI viability assay was used to assess the live and dead cells as previously described (Feng et al., 2014a). With least-square fitting analysis, the regression equation and regression curve of the relationship between percentage of live and dead bacteria as shown in green/red fluorescence ratios was obtained. The regression equation was used to calculate the percentage of live cells in each well of the 96-well plate.

The standard microdilution method was used to determine the MIC of cinnamaldehyde, based on inhibition of visible growth of *B. burgdorferi* by microscopy. 10% cinnamaldehyde DMSO stock was added to *B. burgdorferi* cultures (1 × 10^4^ spirochetes/mL) and was two-fold diluted from 0.5% (= 5 μg/mL) to 0.004% (= 0.04 μg/mL). All experiments were run in triplicate. The

*B. burgdorferi* culture was incubated in 96-well microplate at 33°C for 7 days. Cell proliferation was assessed using the SYBR Green I/PI assay and BZ-X710 All-in-One fluorescence microscope (KEYENCE, Inc.).

### Subculture studies to assess viability of essential oil-treated *B. burgdorferi* organisms

A 7-day old *B. burgdorferi* stationary phase culture (1 ml) was treated with essential oils or control drugs for 7 days in 1.5 ml Eppendorf tubes as described previously (Feng et al., 2015a). After incubation at 33 °C for 7 days without shaking, the cells were collected by centrifugation and rinsed with 1 ml fresh BSK-H medium followed by resuspension in 500 μl fresh BSK-H medium without antibiotics. Then 50 μl of cell suspension was transferred to 1 ml fresh BSK-H medium for subculture at 33 °C for 20 days. Cell proliferation was assessed using SYBR Green I/PI assay and fluorescence microscopy as described above.

## RESULTS

### Evaluation of essential oils for activity against stationary phase *B. burgdorferi.*

In this study, we evaluated another panel of 35 new essential oils at two different concentrations (0.2% and 0.1%) for activity against a 7-day old *B. burgdorferi* stationary phase culture in 96-well plates with control drugs for 7 days. Our previous study discovered cinnamon bark essential oil showed very strong activity against stationary phase *B. burgdorferi* even at 0.05% concentration (Feng et al., 2017). To identify the active components of cinnamon bark essential oil, we also added cinnamaldehyde (CA), the major ingredient of cinnamon bark, in this screen.

Table 1 outlines the activity of the 35 essential oils and CA against the stationary phase *B. burgdorferi.* Using SYBR Green I/PI assay and fluorescence plate reader, 16 essential oils and CA at 0.2% concentration were found to have strong activity against the stationary phase *B. burgdorferi* culture compared to the control antibiotics doxycycline, cefuroxime, and daptomycin (Table 1). As previously described (Feng et al., 2015b), we calculated the residual live cell ratio of microscope images using Image Pro-Plus software, which could eliminate the background autofluorescence. Using fluorescence microscopy, we confirmed the 16 essential oils and CA that could eradicate all live cells as shown by red (dead) aggregated cells (Table 1; Figure 1). At 0.1% concentration, 10 essential oils (garlic, allspice, cumin, palmarosa, myrrh, hedycheim, amyris, thyme white, litsea cubeba, lemon eucalyptus) and CA still exhibited strong activity against the stationary phase *B. burgdorferi* by fluorescence microscope counting after SYBR Green I/PI assay (Table 1; Figure 2). Among them, the most active essential oils were garlic, allspice, cumin, palmarosa, myrrh and hedycheim because of their remarkable activity even at 0.1%, as shown by totally red (dead) cells with SYBR Green I/PI assay and fluorescence microscope tests (Figure 1). CA also showed very strong activity at 0.1% concentration. However, the other 8 essential oils (deep muscle, cornmint, fennel sweet, ho wood, carrot seed, birch, petitgrain and head ease) showed less activity at 0.1% concentration (Table 1, Figure 2). In addition, although birch and litsea cubeba essential oils have autofluorescence, which interfered with the SYBR Green I/PI plate reader assay (showed false high residual viability), they both exhibited strong activity against the stationary phase *B. burgdorferi* as confirmed by SYBR Green I/PI fluorescence microscopy.

**FIG 1.**
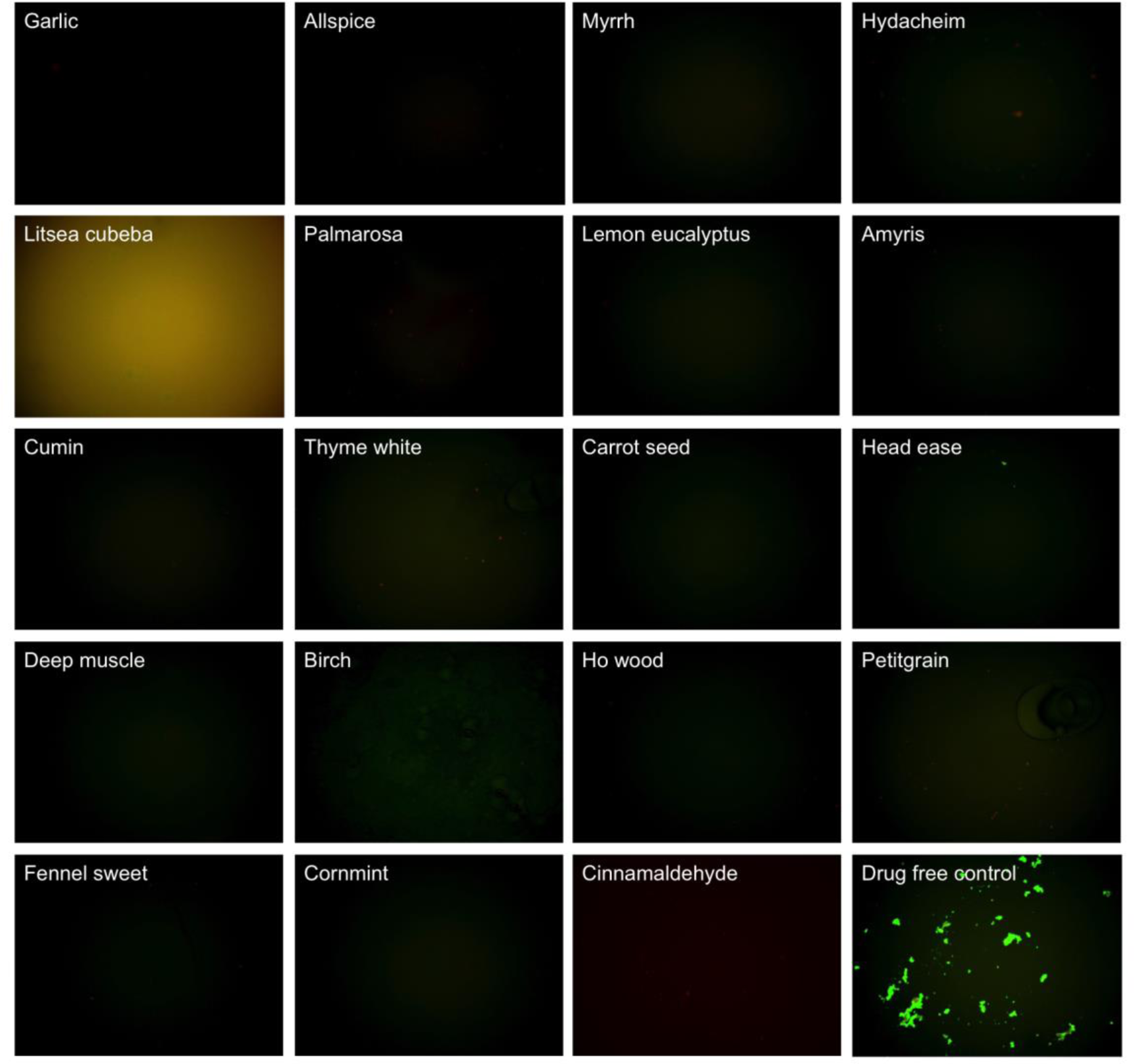
Effect of 0.2% essential oils on the viability of stationary phase *B. burgdorferi.* A 7-day old *B. burgdorferi* stationary phase culture was treated with 0.2% (v/v) essential oils for 7 days followed by staining with SYBR Green I/PI viability assay and fluorescence microscopy.

**FIG 2.**
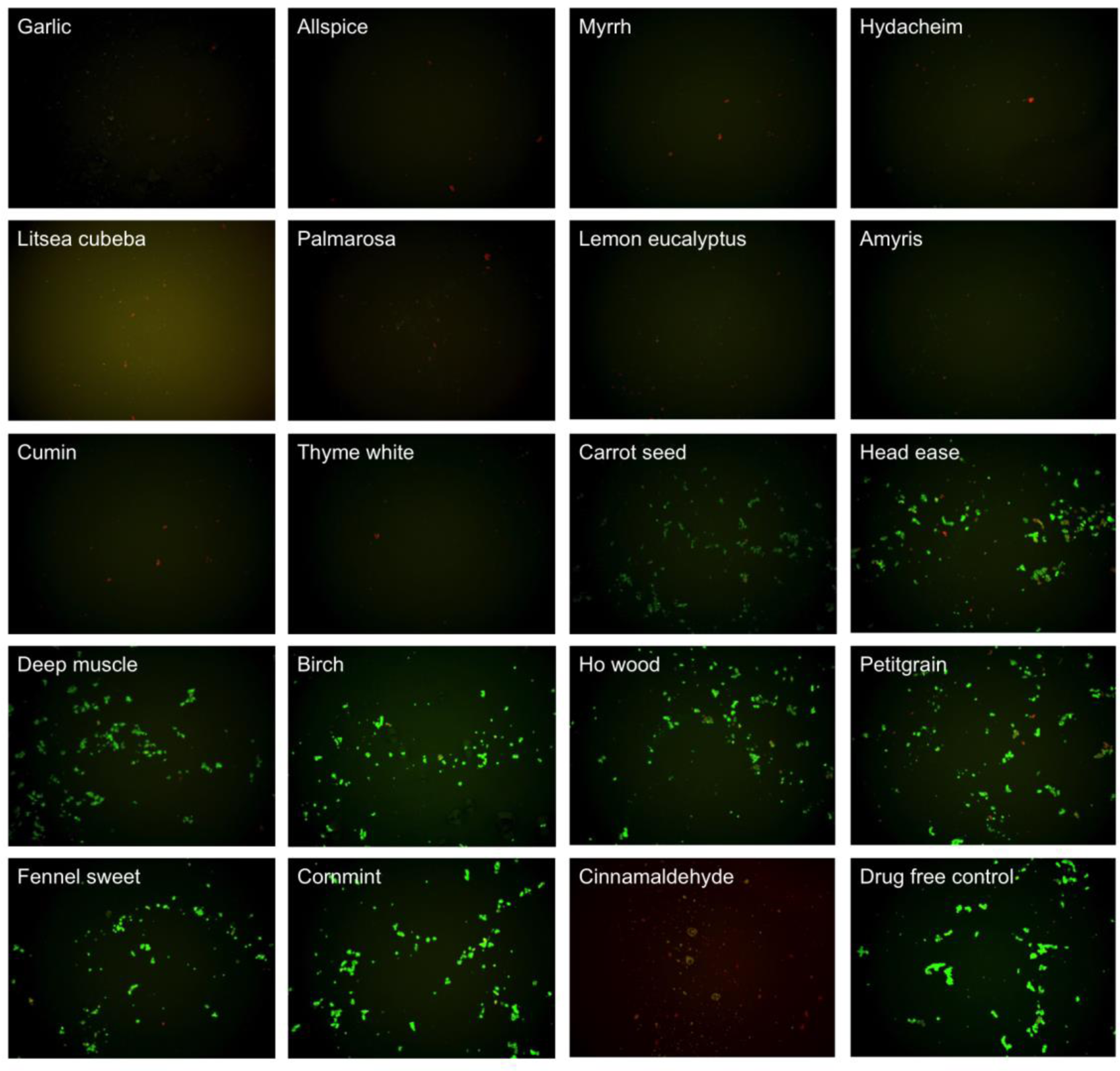
Effect of 0.1% essential oils on the viability of stationary phase *B. burgdorferi.* A 7-day old *B. burgdorferi* stationary phase culture was treated with 0.1% (v/v) essential oils for 7 days followed by staining with SYBR Green I/PI viability assay and fluorescence microscopy.

**Table 1.**
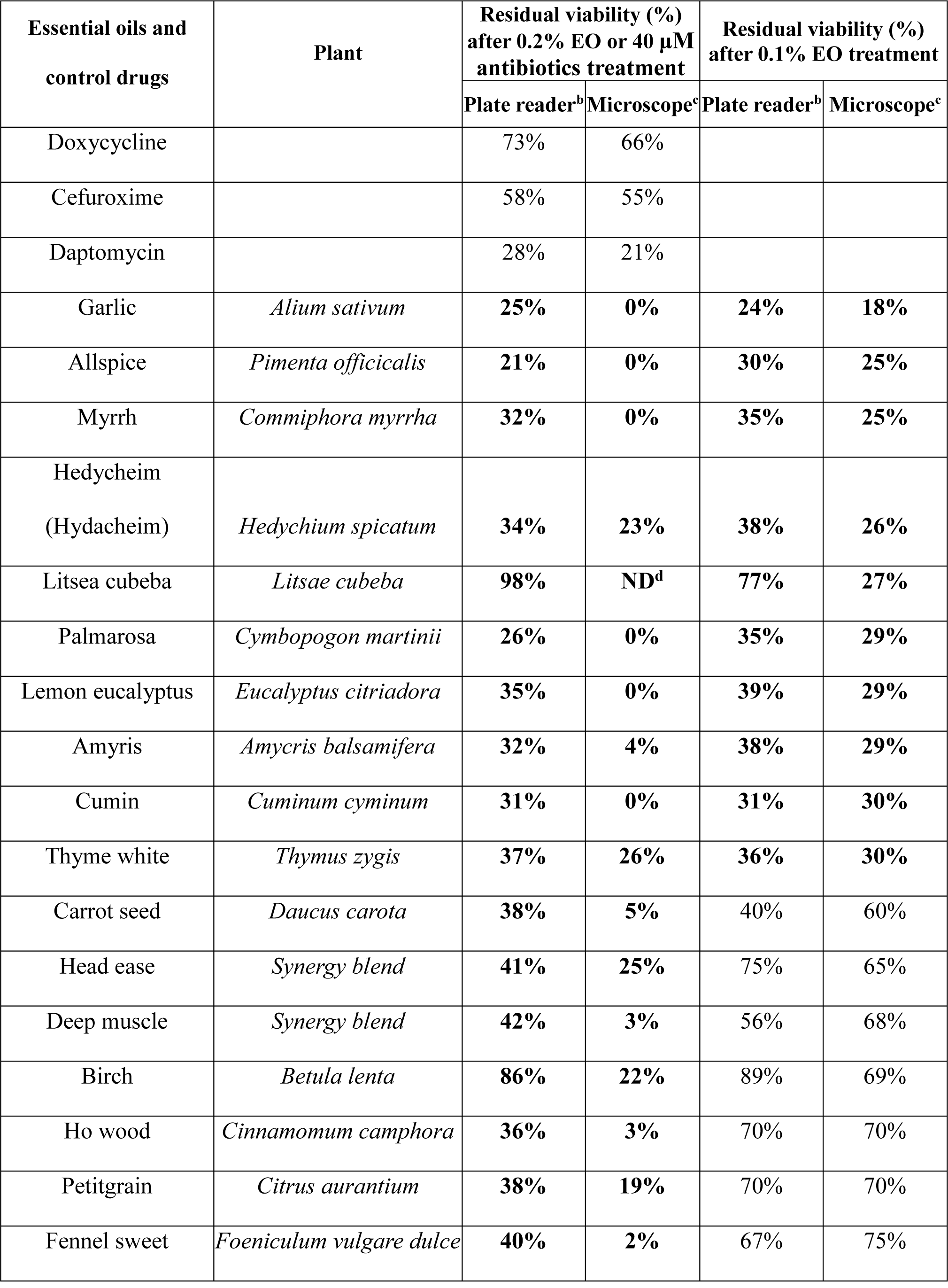
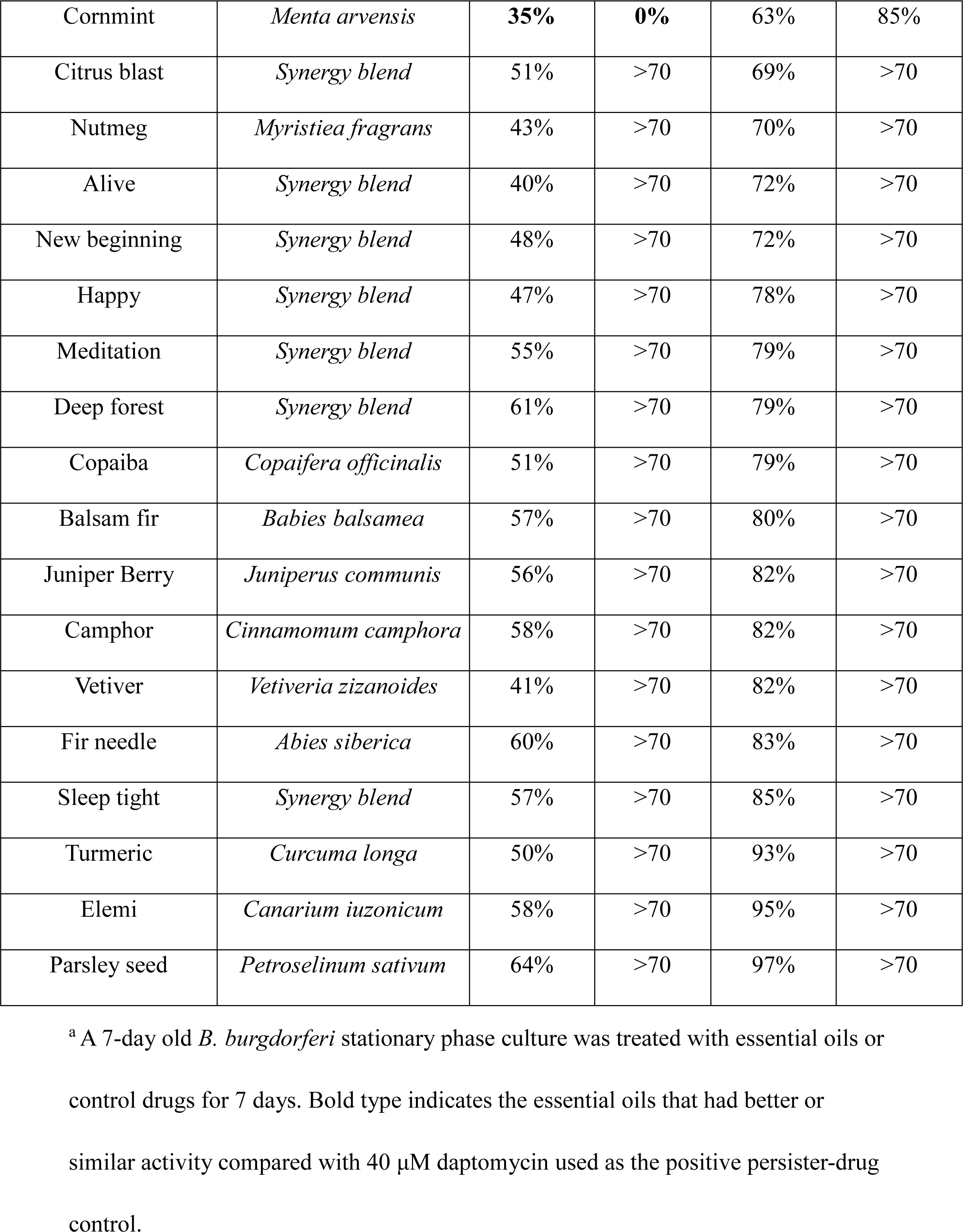
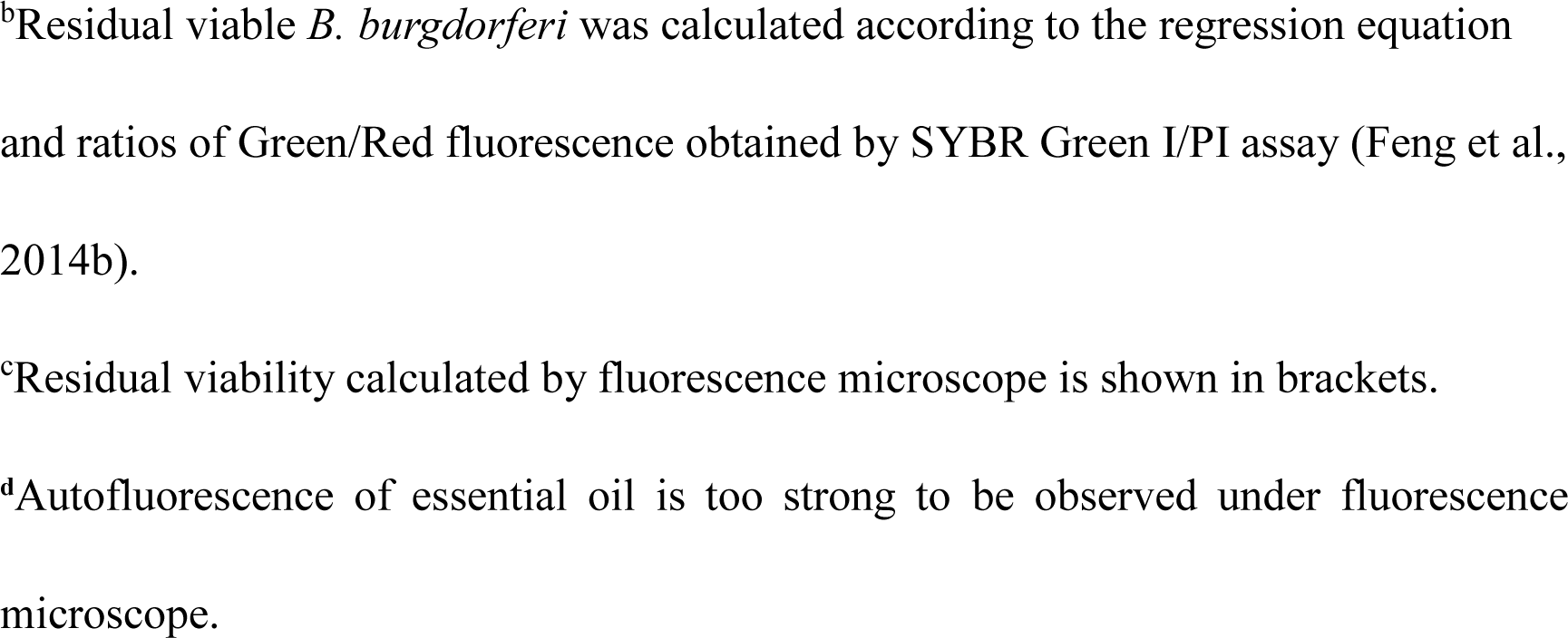
Effect of essential oils on a 7-day old stationary phase *B. burgdorferi* ^a^.

We picked the top 10 essential oils and CA (residual viability lower 60%) to further compare the activity of these active essential oils and determine whether they could eradicate stationary phase *B. burgdorferi* cultures which harbor large numbers of persisters at lower essential oil concentrations (0.1% and 0.05%). We did the confirmation tests with 1 mL stationary phase *B. burgdorferi* in 1.5 mL Eppendorf tubes. At 0.1% concentration the tube tests confirmed the active hits from the previous 96-well plate screen, although the activity of all essential oils decreased slightly in the tube tests compared to the 96-well plate tests (Table 2, Figure 3). At a very low 0.05% concentration, we noticed that garlic, allspice, palmarosa and CA still exhibited strong activity against the stationary phase *B. burgdorferi* as shown by few residual green aggregated cells (Table 2, Figure 3). Meanwhile, we also found CA showed strong activity against replicating *B. burgdorferi*, with an MIC of 0.02% (equal to 0.2 μg/mL).

**FIG 3.**
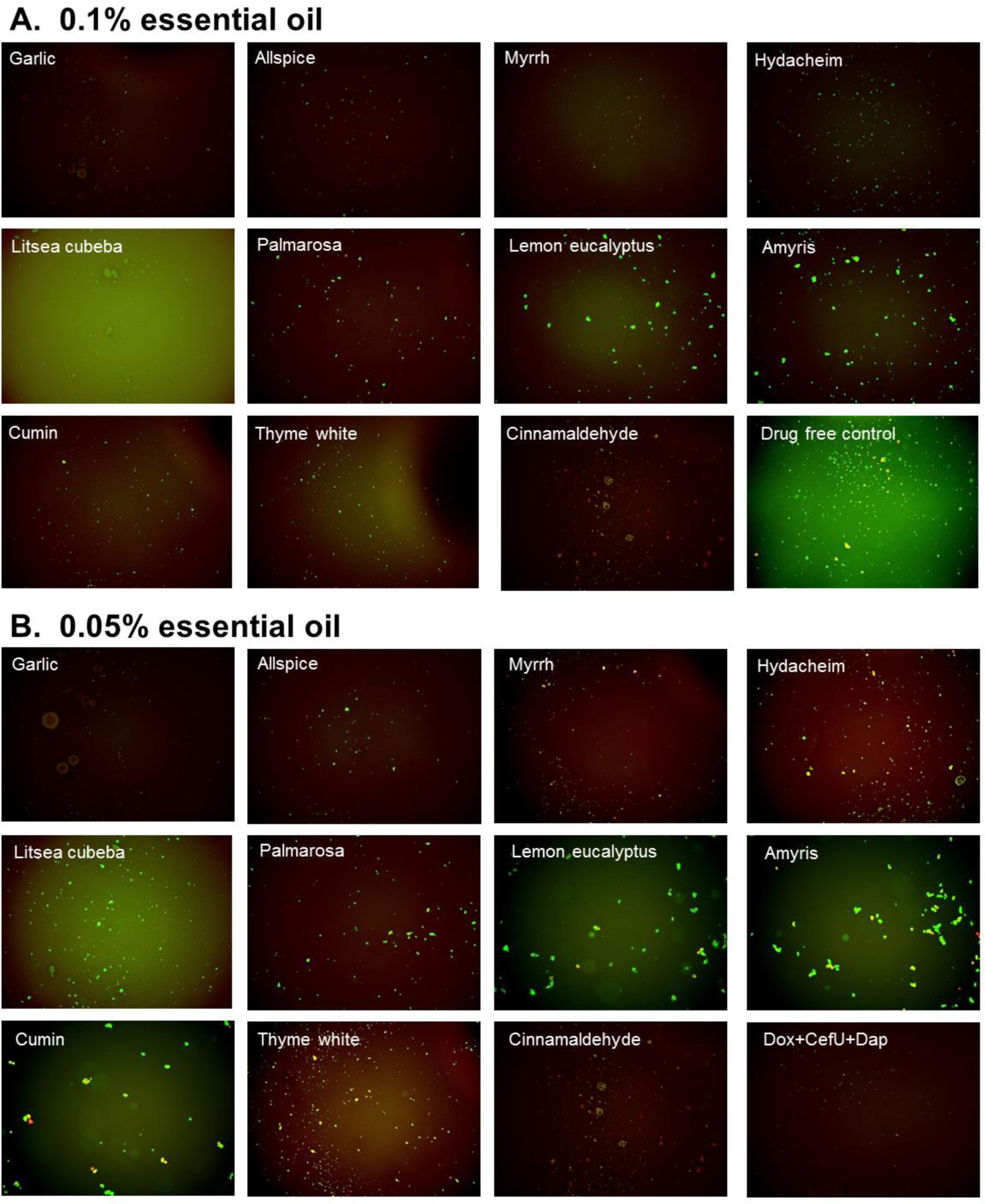
Effect of active essential oils on stationary phase *B. burgdorferi.* A 1 mL *B. burgdorferi* stationary phase culture (7-day old) was treated with 0.1% (A) or 0.05% (B) essential oils (labeled on the image) in 1.5 ml Eppendorf tubes for 7 days followed by staining with SYBR Green I/PI viability assay and fluorescence microscopy.

**Table 2.** Comparison of top 10 essential oil activities against stationary phase *B. hurgdorferi* with 0.1% and 0.05% (v/v) treatment and subculture^a^.

|  | Residual viability after 0.1%<br>Essential oil treatment |  | Residual viability after 0.05%<br>Essential oil treatment |  |
| --- | --- | --- | --- | --- |
|  | Treatment<sup>b</sup> | Subculture<sup>c</sup> | Treatment<sup>b</sup> | Subculture<sup>c</sup> |
| Drug free control | 93% | + | 93% | + |
| Daptomycin+Doxycycline+Cefuroxime<sup>d</sup> | 18%<sup>d</sup> | -<sup>d</sup> | N/A | N/A |
| Garlic | 30% | - | 33% | - |
| Allspice | 34% | - | 48% | + |
| Myrrh | 42% | - | 41% | + |
| Hedychium | 44% | - | 61% | + |
| Litsea cubeba | 68% | - | 69% | + |
| Palmarosa | 39% | + | 67% | + |
| Lemon eucalyptus | 46% | - | 79% | + |
| Amyris | 48% | + | 71% | + |
| Cumin | 42% | + | 60% | + |
| Thyme white | 40% | + | 76% | + |
| Cinnamaldehyde | 34% | - | 56% | - |

<sup>a</sup> A 7-day old stationary phase *B. burgdorferi* was treated with 0.05% or 0.1 % essential oils for 7 days when the viability of the residual organisms was assessed by subculture.
<sup>b</sup>Residual viable percentage of *B. burgdorferi* was calculated according to the regression equation and ratio of Green/Red fluorescence obtained by SYBR Green I/PI assay as described (Feng et al., 2014b). Viabilities are the average of three replicates.
<sup>c</sup> “+” indicates growth in subculture; “-” indicates no growth in subculture.
<sup>d</sup>Activity was tested with 5 µg/mL of each antibiotic in combination.

**Subculture studies to evaluate the activity of essential oils against stationary phase *B. burgdorferi.*** To validate the activity of the essential oils in eradicating stationary phase *B. burgdorferi*, we performed subculture studies after removal of the drugs by washing followed by incubation in fresh BSK medium as previously described (Feng et al., 2015a). We picked the top 10 active essential oils (garlic, allspice, myrrh, hedycheim, litsea cubeba, palmarosa, lemon eucalyptus, amyris, cumin and thyme white) to further confirm whether they could eradicate the stationary phase *B. burgdorferi* cells at 0.1% or 0.05% concentration by subculture experiments after the essential oil exposure (Table 2). At 0.1% concentration, we did not find any regrowth in samples of the top 5 hits including garlic, allspice, myrrh, hedycheim, and litsea cubeba (Table 2, Figure 4A). However, palmarosa, lemon eucalyptus, amyris, cumin, thyme white could not eradicate the stationary phase *B. burgdorferi* as many spirochetes were visible after 21-day subculture (Figure 4A). The subculture study also confirmed the strong activity of cinnamaldehyde by showing no spirochete regrowth in the 0.1% CA treated samples. At 0.05% concentration, we observed no spirochetal regrowth after 21-day subculture in the garlic essential oil treated samples (Figure 4B), which indicates that garlic essential oil could completely kill all *B. burgdorferi* forms even at 0.05% concentration. On the other hand, the other four active essential oils (allspice, myrrh, hedycheim and litsea cubeba) could not sterilize the *B. burgdorferi* stationary phase culture at 0.05%, since they all had visible spirochetes growing after 21-day subculture (Figure 4B). As the previous subculture result of cinnamon bark essential oil (Feng et al., 2017), 0.05% cinnamaldehyde sterilized the *B. burgdorferi* stationary phase culture as shown by no regrowth after 21-day subculture (Figure 4B), which suggests that the active component of cinnamon bark essential oil is attributable to cinnamaldehyde.

**FIG 4.**
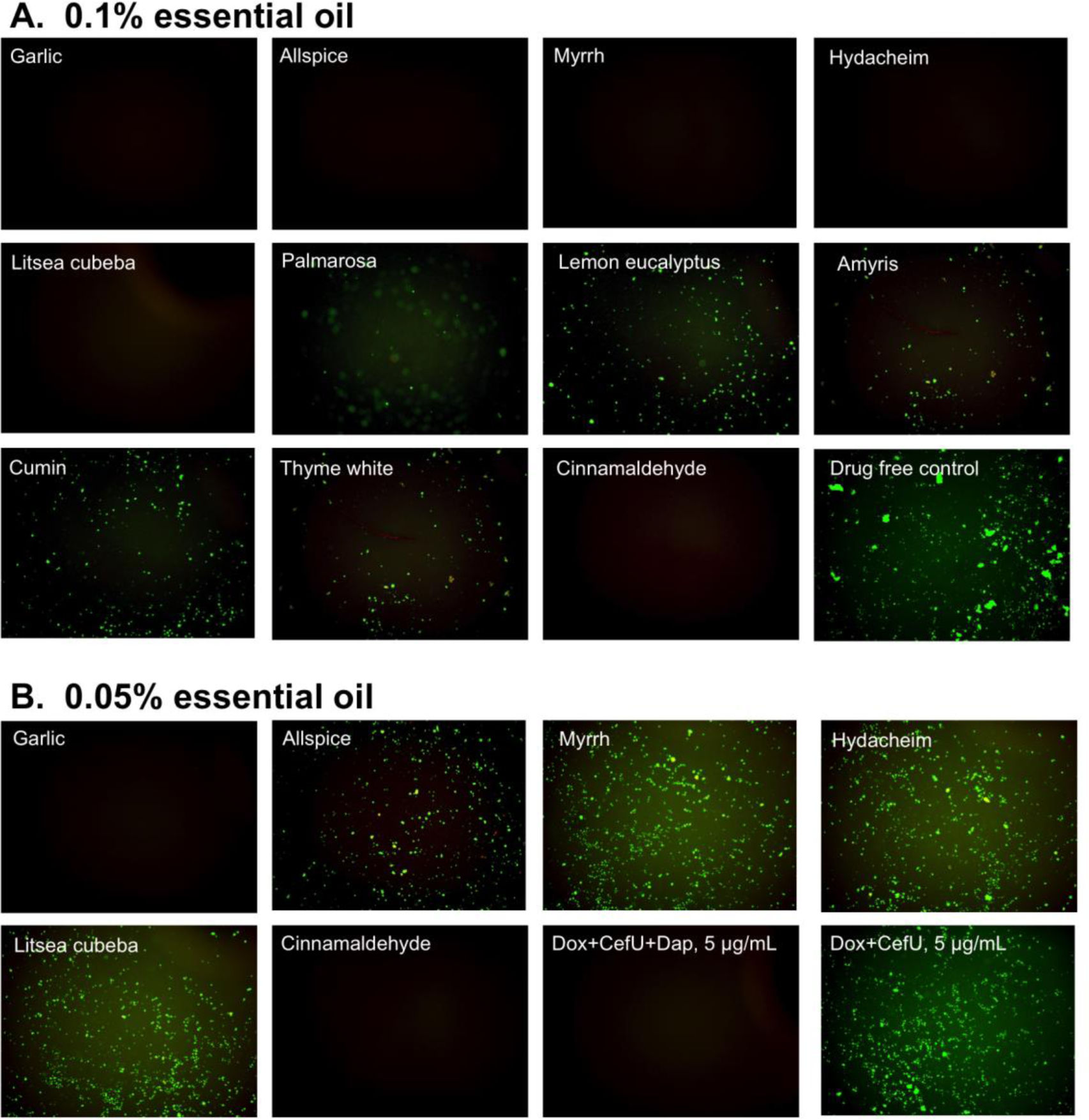
Subculture of *B. burgdorferi* after treatment with essential oils. A *B. burgdorferi* stationary phase culture (7-day old) was treated with the indicated essential oils at 0.1% (A) or 0.05% (B) for 7 days followed by washing and resuspension in fresh BSK-H medium and subculture for 21 days. The viability of the subculture was examined by SYBR Green I/PI stain and fluorescence microscopy.

## DISCUSSION

We recently found many essential oils have better activity against stationary phase *B. burgdorferi* than the current clinically used antibiotics (Feng et al., 2017). Here, we screened another panel of 35 new essential oils using the stationary phase *B. burgdorferi* culture persister model (Feng et al., 2015a). In our previous study, we found 23 essential oils that had strong activity at 1% concentration but only five of them showed good activity at lower 0.25% concentration (Feng et al., 2017). To identify the essential oils which have activity against *B. burgdorferi* persisters at low concentrations, we performed the screen at lower 0.2% and 0.1% concentrations in this study. Although some essential oils like litsea cubeba oil showed high autofluorescence which severely interfered with the SYBR Green I/PI assay (Table 1, Figure 1), using lower concentration (0.1%) and fluorescence microscopy, we were able to verify the results of the SYBR Green I/PI assay. As shown in our previous study, 40 μM daptomycin used as a positive control drug could eradicate *B. burgdorferi* stationary phase cells (Feng et al., 2017), here we identified 18 essential oils that are more active than 40 μM daptomycin at 0.2% concentration, from which 10 essential oils stand out as having a remarkable activity even at 0.1% concentration (Table 1). Among them garlic essential oil exhibited the best activity as shown by the lowest residual viability of *B. burgdorferi* at 0.1%. In the subsequent comparison studies the garlic essential oil highlighted itself as showing a sterilizing activity even at lower concentration of 0.05% because no borrelia cells grew up in the subculture study (Table 2). Garlic as a common spice has been used throughout history as an antimicrobial, and a variety of garlic supplements have been commercialized as tablets and capsules. The antibacterial activity of garlic was described by ancient Chinese and in more recent times by Louis Pasteur in 1858. Although garlic and allicin, an antibacterial compound from garlic, are shown to have antibacterial activity against multiple bacterial species (Lawson, 1998;Petrovska and Cekovska, 2010;Marchese et al., 2016), it has not been well studied on *B. burgdorferi*, especially the nongrowing stationary phase organism, despite its anecdotal clinical use by some patients with Lyme disease (http://www.natural-homeremedies.com/blog/best-home-remedies-for-lyme-disease/; http://lymebook.com/blog/supplements/garlic-allimax-allimed-alli-c-allicin/). In this study, we identified garlic essential oil as having highly potent activity against stationary phase *B. burgdorferi*, and its activity is equivalent to that of oregano and cinnamon bark essential oils, the two most active essential oils against *B. burgdorferi* we identified in our previous study (Feng et al., 2017).

Additionally, we found four other essential oils, allspice, myrrh, hedycheim and litsea cubeba that showed excellent activity against stationary phase *B. burgdorferi*, though the extracts or essential oils of these four plants were reported to possess antibacterial activity on other bacteria. Allspice is a commonly used flavoring agent in food processing and is known to have antibacterial activities on many organisms (Shelef et al., 1980). Myrrh as a traditional medicine has been used since ancient times. In modern times, myrrh is used as an antiseptic in topical and toothpaste (Lisa et al., 2017). It has been shown that some components of myrrh possess *in vitro* bactericidal and fungicidal activity against multiple pathogenic bacteria including *E. coli, S. aureus, P. aeruginosa and C. albicans* (Dolara et al., 2000). Hedycheim (alternate name: hydacheim) essential oil is extracted from the flower of *Hedychium spicatum* plant which is commonly known as the ginger lily plant. The extract of *H. spicatum* is reported to have antimicrobial activity against many bacteria including *Shigella boydii, E. coli, S. aureus, P. aeruginosa* and *K. pneumoniae* (Ritu and Avijit, 2017). Litsea cubeba is also used in traditional Chinese medicine for a long time. Its composition is also reported to exhibit antibacterial activity on *B. subtilis, E. coli, E. faecalis, S. aureus, P. aeruginosa* and *M. albicans* (Wang and Liu, 2010). Based on these studies and application of allspice, myrrh, hedycheim and litsea cubeba, effective regimens may be developed to fight against Lyme disease in the future.

Although these essential oils have high activity against *B. burgdorferi* stationary phase cells *in vitro*, their activity *in vivo* is unknown at this time. In future studies, we will identify the active ingredients of the active essential oils and confirm their activity against growing and nongrowing *B. burgdorferi.* We also need to assess the pharmacokinetic profile of the active compounds in these active essential oils and determine their effective dosage and toxicity *in* vivo. In our previous study we found the cinnamon bark essential oil showed excellent activity against stationary phase *B. burgdorferi* (Feng et al., 2017), here we found CA is an active component of cinnamon bark essential oil. CA could eradicate the stationary phase *B. burgdorferi* at 0.05% concentration as no regrowth occurred in subculture (Table 2). This indicates CA possess similar activity against stationary phase *B. burgdorferi* as carvacrol, which is the one of the most active compounds against non-growing *B. burgdorferi* we identified from natural products (Feng et al., 2017). Furthermore, we also found that CA is very active against growing *B. burgdorferi* with an MIC of 0.2 μg/mL. The antibacterial activity of CA was also reported on some other bacteria. The mechanism of the antibacterial activity of CA has been studied on different microorganisms, which suggests its antibacterial action is mainly through interaction with the cell membrane (Friedman, 2017). CA as a common favoring agent in food processing is also used as food preservative to protect animal feeds and human food from pathogenic bacteria (Friedman, 2017). CA is considered as a safe compound for mammals, as the acute toxicity study showed the median lethal dose LD50 of CA is 1850±37 mg/kg in a rat model (Subash Babu et al., 2007). These findings suggest CA could be a good drug candidate for further evaluation against *B. burgdorferi* in future studies. Appropriate animal studies are necessary to confirm the safety and activity of CA and other active essential oils in animal models before human studies.

In summary, we identified some additional essential oils that have strong activity against stationary phase *B. burgdorferi.* The most active essential oils include garlic, allspice, myrrh, hedycheim and litsea cubeba. Among them garlic oil could completely eradicate stationary phase *B. burgdorferi* with no regrowth at 0.05%, and the others could reach the same activity at 0.1%. Additionally, cinnamaldehyde is identified to be an active ingredient of cinnamon bark with very strong activity against *B. burgdorferi* stationary phase cells. Future studies are needed to identify the active components in the candidate essential oils, determine their safety and efficacy against *B. burgdorferi* in animal models before human trials.

## ACKNOWLEDGMENTS

We acknowledge the support by Global Lyme Alliance, LivLyme Foundation, NatCapLyme, Lyme Disease Association, and Einstein-Sim Family Charitable Fund. YZ was supported in part by NIH grants AI099512 and AI108535.

